# The effect of *Septoria glycines* and fungicide application on the soybean phyllosphere mycobiome

**DOI:** 10.1101/2022.03.23.485325

**Authors:** Heng-An Lin, Santiago X. Mideros

## Abstract

Septoria brown spot (caused by *Septoria glycines*) is the most prevalent soybean disease in Illinois. It is common to use a foliar fungicide to control Septoria brown spot and other foliar diseases. The effects of fungicide on non-target organisms in the phyllosphere are unknown. To study the effect of *S. glycines* and fungicide application on the soybean phyllosphere mycobiome we sequenced full-length ITS and partial LSU region using oxford nanopore technologies. Sequencing produced 3,342 operational taxonomic units (OTUs). The richness of the fungal community significantly increased with the developmental stage. The soybean lines significantly affected the mycobiome diversity at the early developmental stage but not at the reproductive stages. *S. glycines* did not significantly affect the alpha diversity but some significant changes were observed for the beta diversity. At the R5 stage, fungicide application significantly changed the composition of the fungal community. The fungicide treatment significantly decreased the proportion of several fungal reads, but it increased the proportion of *Septoria*. The core microbiome in soybean leaves was composed of *Gibberella, Alternaria, Didymella, Cladosporium, Plectosphaerella, Colletotrichum*, and *Bipolaris*. Network analysis identified significant interactions between Septoria reads and Diaporthe, Bipolaris and two other taxonomic units. In this study, we set *Septoria* as the target organism and demonstrated that metabarcoding could be a tool to quantify the effect of multiple treatments on the fungal phyllosphere community. Better understanding the dynamics of the phyllosphere microbiome is necessary to start untangling late-season diseases of soybean.

## Introduction

Septoria brown spot, caused by *Septoria glycines* Hemmi, is the most prevalent fungal foliar diseases of soybean (*Glycine max* L. Merrill) in Illinois, USA (Hobbs et al. 2010). The estimated yield loss caused by this disease ranged between 1.7 to 6.9 Mt from 2010 to 2014 in the northern USA and Ontario, Canada (Allen et al. 2017). Although the susceptibility of the disease varies among lines, there are no resistance sources or genes for this disease (Hartman et al. 2015). The fungal pathogen, *S. glycines*, overwinter in the soybean residue by forming pycnidia. The pycnidiaspores produced in the next growing season spread by splashing rain to the seedlings and start the initial epidemics from the low canopy. The typical symptoms of this disease are irregular brown necrotic lesions surrounded by an extensive chlorotic area. The vertical progress of the disease development at the late stages of soybean maturity is positively associated with the yield loss. We previously reported that when the disease symptoms reach up to 30% of the plant height at the R6 stage, it leads to a 10% of estimated yield loss, and if it reaches 80% of the plant height, it leads to a 27% of estimated yield loss (Lin et al. 2020). Septoria brown spot can co-occur with other foliar diseases, such as Cercospora leaf blight and anthracnose, and they are considered a late-season disease complex in some references (Carmona et al. 2011). However, the association between *S. glycines* and other pathogenic or non-pathogenic organisms in the soybean phyllosphere has not been studied.

The collective communities of plant-associated microorganisms are referred to as the plant microbiome (Mendes et al. 2013). The members of the plant microbiome can be beneficial, neutral, or pathogenic microorganisms. All parts of the plants can provide a habitat for microorganisms. Based on where those organisms reside, the microbiome can be subdivided into phyllosphere (the aerial parts of plants), rhizosphere (the narrow zone of soil around the plant root system), and endosphere (inside of the plant) microbiome. The plant microbiome is considered an important factor that can be directly or indirectly related to plant nutrient uptake, productivity, disease resistance, and stress tolerance (Turner et al. 2013; Trivedi et al. 2020; Berendsen et al. 2012; Wagg et al. 2011). The microbiome composition differs between plant species, genotype (Sharaf et al. 2019), and developmental stage (Copeland et al. 2015). The communities can also be affected by different agricultural management, such as tillage, fertilization, and chemical treatment (Wattenburger et al. 2019; Longley et al. 2020; Karlsson et al. 2014; Noel et al. 2021). Understanding the effect of agricultural practices on the microbiome is important for the development of sustainable management strategies.

Several studies have investigated the effect of different factors on the microbiome communities in the aboveground of soybeans. The main factors that drive the phyllosphere microbiome communities in soybeans include sampling location, plant growth stage (Longley et al. 2020; Copeland et al. 2015), and agricultural systems (conventional, no-till, or organic farming system) (Longley et al. 2020). Some microbes were abundant in leaf and stem across all management systems, such as *Alternaria*, but some microbes were associated with specific management systems. For example, *Phoma* and *Davidiella* were more abundant in the aboveground samples from the organic farming system (Longley et al. 2020). The application of foliar fungicides is another critical factor that shapes the phyllosphere fungal microbiome (mycobiome). While the desired effect of chemicals on the target organisms was important, more studies start to investigate the impact of those chemicals on the non-target organisms since they might act as competitors or antagonists in the community. In wheat, fungicide application had a moderate but significant effect on the fungal community composition in the phyllosphere (Karlsson et al. 2014). In Karlsson et al (2014), the fungicide application reduced the evenness of the communities. However, the relative abundance of common pathogens in wheat, such as *Mycosphaerella graminicola* and *Parastagonospora nodorum* were not significantly different between control and fungicide treated samples. The relative abundance of those pathogens tends to have higher variation in the fungicide treated samples. In soybeans, Batzer and Mueller (2020) reported the effect of foliar fungicide on soybean endophytes using a cultural-dependent method. They reported that spraying the fungicide decreased the amount of *Alternaria* but increased the proportion of *Diaporthe*, which is the causal agent of soybean pot and stem blight. Although using a cultural-dependent method ensured that the isolates were truly endophytes, they also state that a low diversity of organisms was detected using this method. Using sequencing technologies can overcome the limitation mentioned above and better understand the impact on overall communities in the phyllosphere.

Amplicon sequencing has become a widely used approach for studying the plant microbiome (Lucaciu et al. 2019). The fungal organisms are typically characterized by the internal transcribed spacer (ITS) region, which comprises ITS1 region, 5.8S rRNA gene, and ITS2 region (Raja et al. 2017). The analysis of the mycobiome has mainly been based on sequencing ITS1 or ITS2 regions using second-generation sequencing platforms (e.g., Illumina MiSeq). However, the short amplicons (250 to 450 bp) limit the resolution of phylogenetic information. The entire length of the ITS region can vary from 300 to 1200 bp, with an average length of 600 bp (Heeger et al. 2018; Op De Beeck et al. 2014). Several studies have used third-generation sequencing technologies that produce longer amplicons, showing that using full-length ITS regions or even longer regions (SSU and LSU) for fungal metabarcoding can significantly improve the taxonomic resolution (Heeger et al. 2018; Tedersoo et al 2018).

Unlike the rhizosphere microbiome, we have little information about the phyllosphere microbiome in most plants (Berg et al. 2017). In this study, our goals were: 1) to characterize the mycobiome in soybean phyllosphere using a long-read sequencing platform (Oxford Nanopore), 2) to study the effect of *S. glycines* and fungicide application on the soybean phyllosphere mycobiome, and 3) to study the differences between fungal communities at different developmental stages and soybean lines. We expected that the composition of the fungal communities will change not only by growth stages but also change after fungicide application. Septoria brown spot of soybeans is only one of many fungal species that affect soybeans at maturity, but it is the most commonly observed foliar pathogen in soybeans in Illinois. Understanding the association of *S. glycines* with other foliar fungal organisms and the effect of foliar fungicide on them is necessary to start untangling the late-season disease complex and provide information for better control of the disease.

## Materials and Methods

### Field trial and sample collection

A field trial was conducted in 2018 at the Crop Sciences Research and Education Center near Urbana (N40.070431°, W88.218929°) in Illinois, USA. The field was on corn-soybean rotation and fertilized according to local practices. The experimental design followed a split-plot design with four treatments as the main plots and three soybean lines (Williams, LD12-8677, and LD13-14071R2) as the sub-plots. Soybeans were planted on 30 April at 348,480 seeds hec^-1^, and each sub-plot consisted of four 5.2 m long rows with 76.2 cm alleys.

The treatments consisted of 1) Inoc: inoculated with *Septoria glycines* at V3 stage (Fehr et al. 1971); 2) Inoc/Fun: inoculated with *Septoria glycines* and received fungicide application at R3 stage; 3) NIC/Fun: not inoculated and received fungicide application at R3 stage, and (4) NIC/No_Fun: not inoculated nor fungicide application control. Each treatment had three replicates. Three *S. glycines* isolates (16S006, 16S012, and R3216) that stably sporulated were chosen for inoculation. The culture conditions of the fungal isolates, preparation of the inoculum, and inoculation procedure followed the description in Lin et al. (2020). The center two rows of the assigned plots were inoculated with a suspension of 10^6^ spores mL^-1^ at the V3 stage (5 June). A second inoculation was applied in those plots at the R2 stage (3 July) to increase the disease severity. Fluxapyroxad and pyraclostrobin (Priaxor, BASF) was sprayed at the R3 stage (17 July) at the assigned plots with a CO_2_-pressurized backpack sprayer equipped with a 0.48 m 601B-SST-four nozzle light weight boom (R&D Sprayers Bellspray, Inc, USA) equipped with four TJ60-11008 (50) nozzles (TeeJet Technology, USA). Ten plants were randomly collected from each subplot at three timepoints, (1) One week after inoculation treatment (V4 stage, collected on 11 June, 13 June, and 14 June), (2) One week before fungicide application (R3 stage, collected on 10 July, 11 July, and 12 July), and (3) One week after fungicide application (R5 stage, collected on 23 July, 24 July, and 25 July) stage. The samples were collected by replicate in the morning and processed within the same day. Percentage of necrotic area, chlorotic area, vertical progress, and defoliation rate were evaluated for each plant (Lin et al. 2020). After disease evaluation, all leaves were collected from each plant. In total, 108 pooled leaf samples were lyophilized and stored at -20°C for DNA extraction.

### DNA extraction

The lyophilized leaf samples were submerged in liquid nitrogen and then mixed in a laboratory blender (Waring Commercial, USA). A total of 100 mg of mixed leaf tissue from each sample were placed in a 2 mL FastPrep^®^ tube containing a 6.35 mm ceramic bead (MP BIOMEDICALS, USA) for homogenization and DNA extraction. The DNA was extracted following a modified CTAB method as described in Lin and Mideros (2021). The extracted DNA was analyzed on a NanoDrop spectrophotometer (Thermo Scientific™, USA).

### Library preparation and sequencing

Ninety-six out of 108 samples with good quality of DNA were sent to Roy J. Carver biotechnology center for library preparation and sequencing (Supplementary Table 1). The sequenced samples consisted of three replicates for each treatment collected at the V4 and R3 stage, and two replicates for each treatment at the R5 stage. The amplicons were produced with the Fluidigm Access Array system (Fluidigm, USA) using primers ITS1F (5’-CTTGGTCATTTAGAGGAAGTAA-3’) and LR6 (5’- CGCCAGTTCTGCTTACC-3’) (Walder et al. 2017). The ITS1F/LR6 primer set targeted the fungal internal transcribed spacer (ITS) and a partial region of the large subunit (LSU) of the rDNA. The pooled amplicons were converted into an Oxford Nanopore library with the SQK-LSK109 library kit (Oxford Nanopore, UK). The library was sequenced on a SpotON R9.4.1 FLO-MIN106 flow cell for 72 hr, using a GridIONx5 sequencer. Base-calling of the fastq files was performed with Guppy 3.0.3 (https://community.nanoporetech.com).

### Bioinformatics analysis

The filtering of reads with average Q scores < 7 was done with NanoFilt (De Coster et al. 2018). Demultiplexing and library adaptor trimming were done with Porechops 0.2.3 (Wick et al. 2018). The reads that contained adaptors in the middle of the sequence were removed using Porechop 0.2.3 (Wick et al. 2018). Reads were then filtered by length using Nanofilt (De Coster et al. 2018), and the reads with length between 250 to 2000 bp were retained for further analysis. Transformation of the files from fastq to fasta format was performed using FASTX-Toolkit 0.0.14 (Hannon 2010). Clustering into operational taxonomic units (OTUs) for each sample was conducted using VSEARCH 2.4.3 (Rognes et al. 2016). First, the reads were pre-clustered within each sample at 95% similarity. Then a second clustering was conducted in the merged files at 95%, 92%, 90%, 88%, and 86% similarity. The optimal clustering criteria was determined to be 92% because it was the highest similarity that produced an acceptable percentage of singletons (Supplementary Table 2). Singletons and doubletons were filtered out (Allen et al. 2016). The results are called here the OTU frequency table and was generated using a custom bash script (frequencytable.sh).

Multiple sequence alignment was performed within each OTU, and a consensus sequence was extracted from each OTU as described by Mafune et al (2020). Because the UNITE v8.2 database provides full-length ITS sequences and the RDP database provides LSU sequences, for each one of our consensus sequences we extracted the ITS and LSU regions using ITSx 1.1.1 (Bengtsson-Palme et al. 2013). To assign the taxonomy to each OTU, the ITS and LSU data sets were aligned using BLAST to the UNITE v8.2 general fasta release fungi database (Nilsson et al. 2018), and to the RDP v2.9.0 database (Cole et al. 2014), respectively. The ITS and LSU sequences were aligned to the database using BLAST+ 2.9.0 (Camacho et al. 2009) with an E-value set as 10^−5^. The classification of each OTU was determined with the lowest E-value, highest bit score, and highest identity (Supplementary Table 3). The results are called here the OTU taxonomy table and was generated from the BLAST results using a custom bash script (featuretable.sh).

### Statistical analysis of disease severity phenotypes

The area under the disease progress curve (AUDPC) values for the percentage of vertical progress (the percent height of the plant with disease symptoms), necrotic area, chlorotic area, and defoliation rate were calculated using the package Agricolae (De Mendiburu 2010) in R 4.0.2 (R Core Team, 2020). A split-plot model was used to evaluate the experiment.

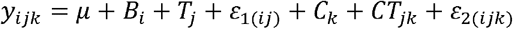

In the model, *y*_*ijk*_ is the AUDPC for each disease severity component corresponding to the *i*^*th*^ block, *j*^*th*^ treatment, and *k*^*th*^ line. *μ* is the grand population mean, *B*_*i*_ is the random block effect, *T*_*j*_ is the fixed treatment effect, *C*_*k*_ is the fixed line effect, *CT*_*jk*_ is the fixed interaction effect between the *j*^*th*^ treatment and *k*^*th*^ line. The *ε*_1(*ij*)_ and *ε*_2(*ijk*)_ are the random error terms 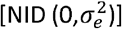. The model was fit using PROC MIXED in SAS v.9.4 (SAS Institute., Cary, NC).

### Statistical analysis of mycobiome diversity

Diversity analyses were performed at both OTU and genus levels. The data was separated into three growth stages (V4, R3, and R5) to study the effect of treatments on the fungal communities. The V4 and R3 stages were used to assess the effect of inoculation treatment. The R5 stage data were used to assess the effect of both inoculation and fungicide application treatments. The differences between three cultivars and growth stages were also analyzed. For the analyses the OTU frequency table, taxonomy table, sample information table, and consensus sequences were imported into R 4.0.2 (R Core Team, 2020) and formatted using *otu_table, tax_table, sample_data, refseq*, and *phyloseq* functions in the phyloseq package (McMurdie and Holmes 2013). The OTU frequency table (the read counts of each OTU in each sample) was used to calculate richness, Shannon index, and Simpson index using the estimate_richness function in phyloseq. The ANOVA analysis for the alpha diversity indexes was conducted in SAS using the same model as for the phenotypes. The Kruskal-Wallis test was conducted in R using the stats package.

The beta diversity analysis was performed using the OTU frequency and OTU taxonomy tables (for the latter there is ITS and LSU data sets). The reads were normalized using the rarefaction method using the rarefy_even_depth function in phyloseq with default settings (Willis 2019). A range of 30 to 70% of OTUs were removed after normalization, depending on the data set (Supplementary Table 4). To generate the weighted Unifrac distance matrix (Lozupone et al. 2011), phylogenetic trees were constructed using the packages phangorn (Schliep 2011) in R. The principal coordinate analysis (PCoA) plots were generated from the weighted Unifrac distance matrix. Multivariate homogeneity of groups dispersions was performed using the betadisper function in R package vegan (Oksanen et al. 2020) to check if the data met the variance assumption for permutational multivariate analysis of variance analysis (PERMANOVA). A PERMANOVA analysis was performed using the adonis function in R package vegan (Oksanen et al. 2020). The OTU differential abundance analysis was performed using DeSeq2 (Love et al. 2014) to identify the specific OTUs at the genus level that were affected by the treatments..

### Analysis for core OTUs

To identify organisms that compose the core phyllosphere mycobiome in soybean we used two data sets: the sequences from the control samples (NIC/No_Fun) so that there was no interference from inoculation or fungicide application, and all the samples. The core mycobiome was defined as OTUs with a prevalence greater than or equal to 0.5 and abundance larger than 1% using package microbiome (Lahti et al. 2012-2017) in R. To confirm the identification, consensus sequences of the core microbes were aligned using BLASTn to the genbank and Unite databases (Supplementary table 5). A split-plot model (same used for the phenotypes) was used to evaluate the experiment for each core OTU. In the model, *y*_*ijk*_ was the relative abundance for each OTU corresponding to the *i*^*th*^ block, *j*^*th*^ treatment, and *k*^*th*^ lines. The other terms were as indicated above for the disease severity phenotypes. The data were transformed as needed to meet the assumptions for ANOVA analysis. The model was fit using PROC MIXED in SAS v.9.4 (SAS Institute., Cary, NC).

### Network analysis

The co-occurrence network analysis was performed using Sparse InversE Covariance Estimation for Ecological Association Inference (SPIEC-EASI) method (Kurtz et al. 2015) in R. The unnormalized data were merged to the genus levels and subsetted based on the sampling stage (V4, R3 and R5) and treatments. The V4 data set include 18 inoculated and 18 no-inoculated samples; R3 data set include 17 inoculated and 17 not-inoculated samples; and R5 data set included 12 samples for each treatment combinations (Inoc/Fun, NIC/Fun, Inoc/No_Fun, and NIC/No_Fun). In each dataset, OTUs with a frequency of less than six were excluded from the analysis (Kerdraon et al. 2019). The SPIEC-EASI pipeline first transformed the OTU frequency data to centered log ratio data. Then, the Meinshausen-Bühlmann’s neighborhood selection method (MB method) was used as the graphical inference model with the minimum lambda ratio set at 10^−2^ and 50 repetitions. The adjacency matrix and centrality metrics (alpha centrality and betweenness centrality) were extracted from the speiec.easi object (Birt and Dennis 2021). The edge stability of each network was extracted with the getOptMerge function in R package SPIEC-EASI. The complexity of the network was defined by the average number of edges (total edges/total nodes) of a network. The networks were visualized and plotted with Cytoscape V. 3.6.1 (Shannon et al. 2003).

### Data availability

All sequences were submitted to NCBI under the Bioproject PRJNA764246, and all scripts were available at https://github.com/henganl2/Microbiome_data

## Results

### Disease development

Typical symptoms of Septoria brown spot were observed at the R3 stage. At this stage, a higher percentage of vertical progress (40%), necrotic area (4-6%), chlorotic area (4-6%), and defoliation rate (20-30%) were observed on the inoculated plots. While the non-inoculated plots had a 20% of vertical progress, 4-6% of necrotic and chlorotic area, and 20% of defoliation rate. At the next evaluation stage (R5), the disease development did not significantly decrease due to fungicide application, but overall, the disease severity on the non-inoculated plots was lower than the inoculated plots (Supplementary Table 6). The ANOVA indicated that the treatments significantly (*p*<0.05) affected the AUDPC of the percentage of vertical progress, chlorotic area, and necrotic area. The soybean lines also significantly affected the AUDPC of the percentage of vertical progress and necrotic area (Supplementary Table 7).

### Sequencing and OTU classification summary

The Oxford Nanopore GridIONx5 sequencer produced a total of 4,354,575 reads, with a mean read length of 1,819 bp in the range of 300 bp and 9,941 bp. Samples had an average 44,183 reads. Two out of 96 samples were discarded from the analysis. One of these samples had only 642 reads and the other completely failed to produce amplicons. The number of filtered reads was 4,184,494, with an average of 44,047±36,773 (SD) reads for each sample (Supplementary Table 1). There were a total number of 1,490,827 unique OTUs identified, and 3,342 OTUs (2,668,341 reads) remained after filtering out the singletons (35%) and doubletons (0.09%) (Supplementary Table 2). We extracted 2,842 ITS sequences and 2,976 LSU sequences out of 3,342 OTUs’ consensus sequences. After the alignment (with BLAST), there were 2840 hits and 2972 hits for ITS and LSU sequences respectively (Supplementary Table 3). For the OTU classification using the ITS sequences, 94.37% were classified as Ascomycota, followed by Basidiomycota (3.98%), Chytridiomycota (0.07%), Glomeromycota (0.18%), Mortierellomycota(0.04%), leaving 1.37% as unclassified. For the OTU classification using the LSU sequences, 95.49% were classified as Ascomycota, followed by Basidiomycota (3.77%), Chytridiomycota (0.20%), and Blastocladiomycota (0.07%), leaving 0.47% as unclassified. Overall, about 99% of ITS and LSU sequences can be classified to a taxonomy level with the identity ranging between 80 to 100% and a mean of 97%. Comparison between the LSU and ITS taxonomies were limited due to the different formats of the databases and inconsistent naming of organisms (for example Fusarium vs Gibberella). Both taxonomies agreed at the Domain and Phylum levels, but the consensus was less than half at the genus level and completely different at the species level.

### Relative abundance (RA) of *Septoria*

At the V4 stage, the relative abundance (RA) of *Septoria* ranged between 0-7% and 0-3% in the inoculated samples and non-inoculated samples, respectively. The RA of *Septoria* was significantly different between inoculated and non-inoculated samples at V4 stage within lines except for cv. LD13-14071R2. The relative abundance of *Septoria* increased at the R3 stage in both inoculated (0-23%) and non-inoculated samples (0-3%). The significant difference between treatments indicated that the inoculation was successful, and the pathogen successfully colonized the soybean leaf tissue after inoculation. A low amount of *Septoria* was also detected in the non-inoculated samples (0-3%), showing that the pathogen was present naturally in the field. At the R5 stage, the lines were merged to calculate the RA since there was no significant difference between them from the diversity analysis. More *Septoria* was identified in the fungicide treated samples. The RA of Septoria was still higher in the inoculated samples compared to the non-inoculated samples. Although the fungicide application did not decrease the RA of *Septoria*, there was a higher variability of *Septoria* between samples (5±6% SD) (Figure 1).

**Figure 1.**
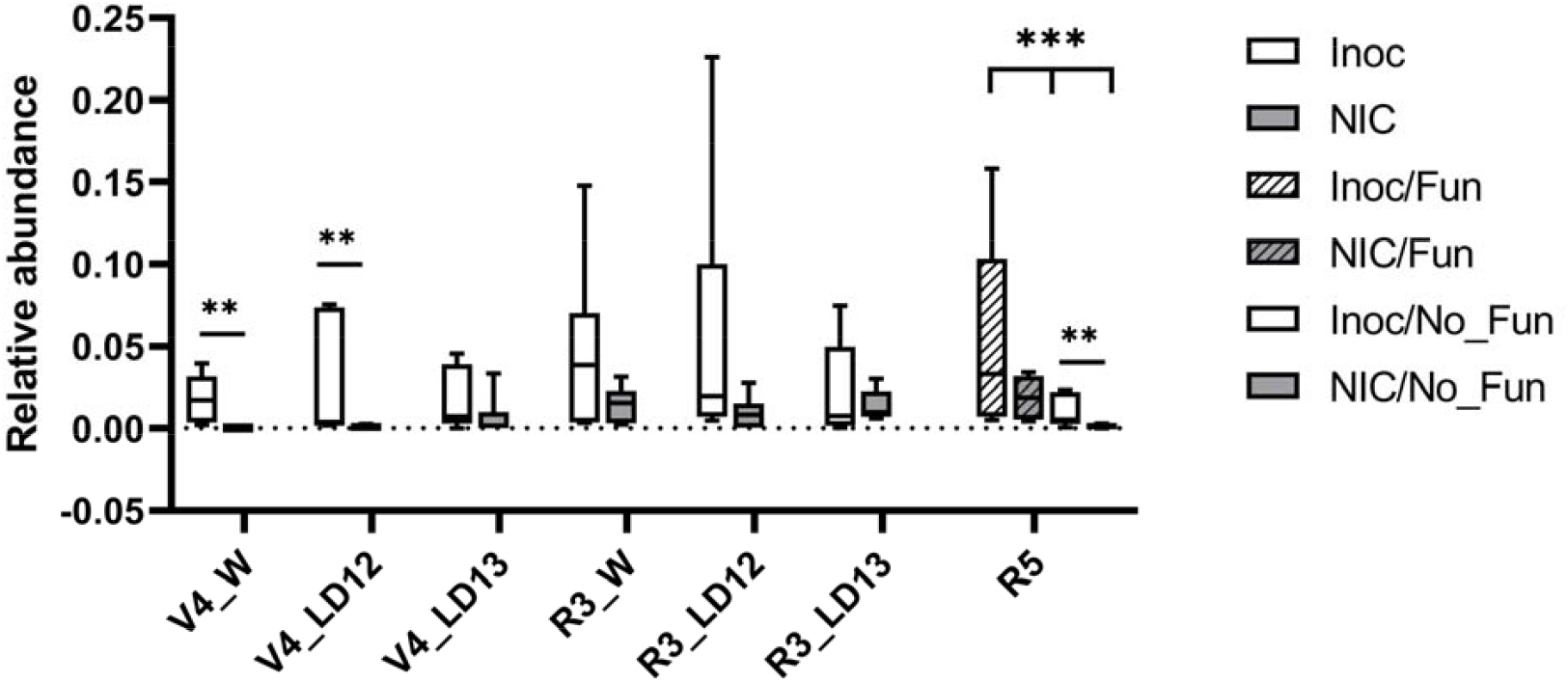
Relative abundance of Septoria at V4, R3, and R5 stage. At V4 and R3 stage, each box consisted of three replicates for W(illiams), LD12(−8677), and LD13(−14071R2). At R5 stage, the lines were merged for analysis since there was no significant difference between lines, and each box consisted of six replicates. The Wilcoxon rank-sum tests were performed for analysis (Inoc vs NIC, Fun vs No_Fun).(*p<0.1, **p<0.05, ***p<0.005)

### Community analysis – Alpha diversity

The alpha diversity was estimated with richness, Shannon index, and Simpson index using the OTU- and genus-level data. The results showed a similar trend between the two data sets, but more significant comparisons were identified from the genus-level data. A significant difference (*p*<0.05) in the ANOVA between lines was only detected at the V4 stage for all diversity indices (Table 1). There was also a significant difference in richness (*p*<0.05) and Simpson index (*p*<0.05) between the three growth stages (Table 2, Figure 2). Richness slightly increased at the R3 and R5 stages. The distribution of the diversity indexes for all three cultivars had the widest range at the R3 stage regardless of the received treatments. At the R5 stage, although the diversity was higher in the fungicide treated samples regardless of inoculation, the fungicide treatment reduced the variation between the replicates. A significant difference (*p*<0.05) between inoculated and non-inoculated treatments was only identified for the richness in cv. Williams at V4 stage (Table 2, Figure 2). This result indicated that the inoculation with *Septoria* did not affect the overall fungal alpha diversity.

**Figure 2.**
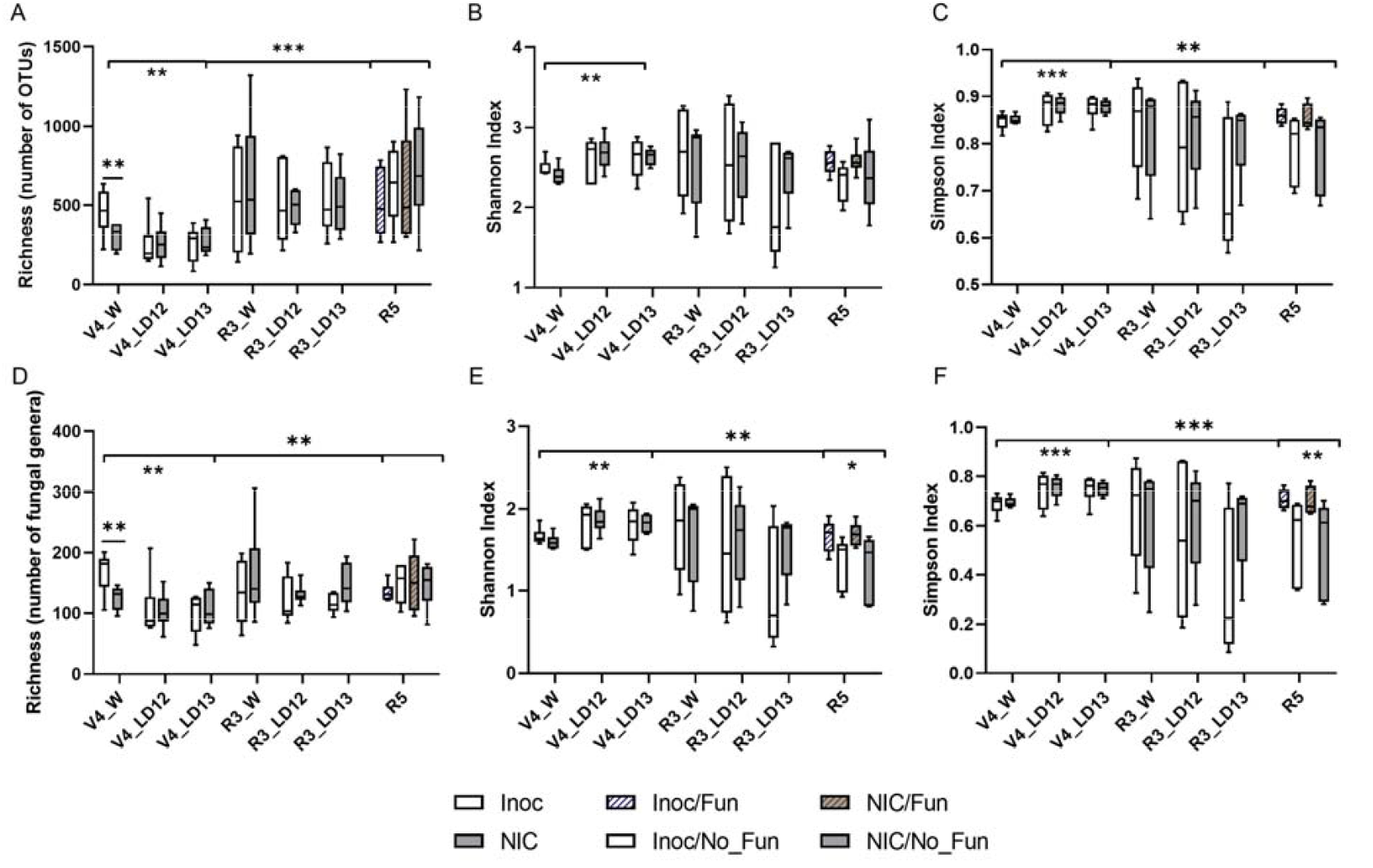
Difference of three alpha diversity indices (Richness, Shannon index, and Simpson index) between treatments, lines, and growth stage using data at (A)-(C) OTU level and (D)-(F) genus level. At V4 and R3 stage, each box consisted of three replicates for W(illiams), LD12(−8677), and LD13(−14071R2). At R5 stage, the lines were merged for analysis since there was no significant difference between lines, and each box consisted of 6 replicates. The Kruskal-Wallis tests were performed for treatments, lines, and growth stage. (*p=0.0506, **p<0.05, ***p<0.005)

**Table 1.** Analysis of variance (ANOVA) to assess the effect of treatments (inoculation and no inoculation and soybean lines) on alpha diversity. Only growth stages with a significant statistically significant value are listed.

| Taxonomy level | Growth stage | Alpha diversity index | Factor | F value | *p* value<sup>2</sup> |
| --- | --- | --- | --- | --- | --- |
| OTUs | V4 | Richness<sup>1</sup> | Inoculation | 0.23 | NS |
|  |  |  | Lines | 4.29 | 0.0237** |
|  |  |  | Inoculation*Lines | 1.5 | NS |
|  |  | Shannon | Inoculation | 0 | NS |
|  |  |  | Lines | 4.37 | 0.0987* |
|  |  |  | Inoculation*Lines | 1.19 | NS |
|  |  | Simpson | Inoculation | 0.13 | NS |
|  |  |  | Lines | 8.74 | 0.0347** |
|  |  |  | Inoculation*Lines | 0.12 | NS |
| Genus | V4 | Richness | Inoculation | 0.43 | NS |
|  |  |  | Lines | 5.31 | 0.0749* |
|  |  |  | Inoculation*Lines | 1.36 | NS |
|  |  | Shannon | Inoculation | 0.01 | NS |
|  |  |  | Lines | 6 | 0.0625* |
|  |  |  | Inoculation*Lines | 0.71 | NS |
|  |  | Simpson | Treatment | 0.15 | NS |
|  |  |  | Lines | 9.35 | 0.031** |
|  |  |  | Inoculation*Lines | 0.12 | NS |

<sup>1</sup>The value was log-transformed to meet the assumption of normality and homogeneity of variance for the ANOVA test.
<sup>2</sup> NS: not significant, \**p*<0.1, \*\**p*<0.05, \*\*\**p*<0.005

**Table 2.** Kruskal Wallis test to assess the effects of treatments^1^ and lines on the alpha diversity index. Only growth stages or line within growth stage with statistically significant results are listed.

| Taxonomy level | Growth stage / Lines | Factor | Alpha diversity index | Chi-squared | *p* value<sup>2</sup> |
| --- | --- | --- | --- | --- | --- |
| OTUs | All lines | Growth stages | Richness | 27.341 | <0.005*** |
|  |  |  | Shannon | 1.8533 | NS |
|  |  |  | Simpson | 9.1614 | 0.010** |
|  | V4 | Lines | Richness | 7.19 | 0.027** |
|  |  |  | Shannon | 7.96 | 0.019** |
|  |  |  | Simpson | 11.28 | 0.004*** |
|  | V4/Williams | Inoculation | Richness | 5.04 | 0.024** |
|  |  |  | Shannon | 2.08 | NS |
|  |  |  | Simpson | 0.03 | NS |
| Genus | All lines | Growth stages | Richness | 7.4599 | 0.024** |
|  |  |  | Shannon | 7.0075 | 0.030** |
|  |  |  | Simpson | 11.562 | 0.003*** |
|  | V4 | Lines | Richness | 9.92 | 0.007** |
|  |  |  | Shannon | 9.54 | 0.008** |
|  |  |  | Simpson | 11.07 | 0.004*** |
|  | V4/Williams | Inoculation | Richness | 4.35 | 0.037** |
|  |  |  | Shannon | 2.08 | NS |
|  |  |  | Simpson | 0.03 | NS |
|  | R5 | Treatments | Richness | 1.29 | NS |
|  |  |  | Shannon | 7.79 | 0.051* |
|  |  |  | Simpson | 8.29 | 0.040** |

<sup>1</sup> Treatments at V4 stage: inoculation and no inoculation; Treatments at R5 stage: inoculation/fungicide, inoculation/no fungicide, no inoculation/fungicide, and no inoculation/no fungicide.
<sup>2</sup> NS: not significant, \**p*<0.1, \*\**p*<0.05, \*\*\**p*<0.005

### Community analysis – Beta diversity

The PERMANOVA test showed that the soybean lines were a significant (*p*=0.001) factor that structured the fungal community at V4 stage at the OTU level (Table 3). A significant effect for the inoculation treatment (*p*=0.03) was only detected at the V4 stage at the OTU level for the ITS data set. While the inoculation treatment explained 1% to 6% of the variation, the lines explained 13% to 37% of variation at the V4 stage. No significant factors were identified from the PERMANOVA at the R3 stage. At the R5 stage, there was no significant difference between lines. The treatments at the R5 stage were only significant at the OTU level and explained 40% of the variation (*p*=0.091), however the there was a significant variation in the multivariate dispersion (not shown). The principal coordinates analysis (PCoA) also suggested that the community clustered by lines at V4 stage and that inoculation had no effect on the community (Figure 3 A-B). No apparent patterns were observed from the PCoA results for the R3 stage data. At the R5 stage, a trend was observed that the application of fungicide had a larger effect than lines and inoculation treatment suggesting that fungicide application shapes the fungal community in the phyllosphere (Figure 3 C-D).

**Figure 3.**
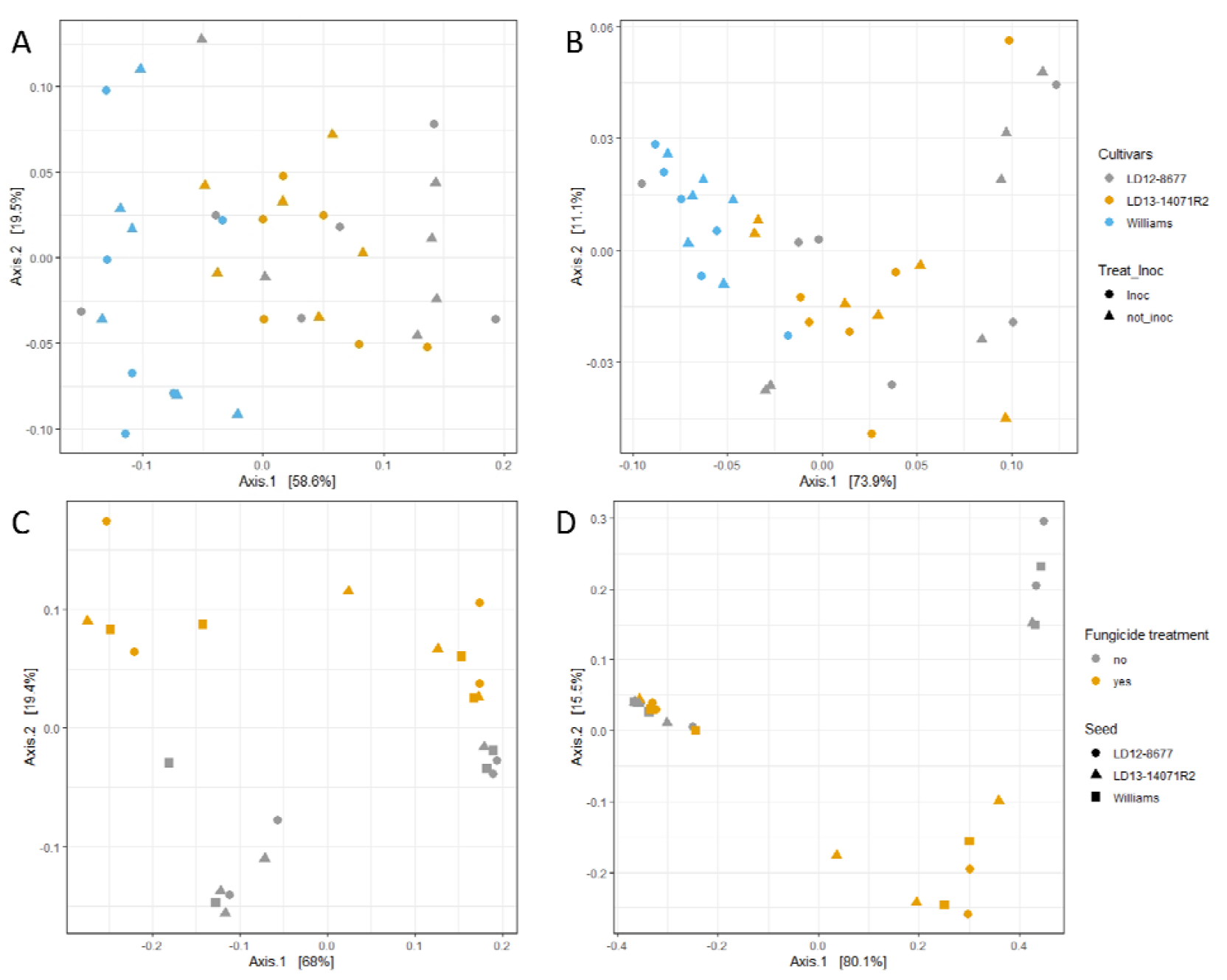
Principal coordinates analysis (PCoA) of weighted unifrac distance matrix. The distance matrices were calculated with the rarefied data (ITS or LSU data set) at the genus level by growth stage. (A) V4, ITS data (B) V4, LSU data (C) R5, ITS data, and (D) R5, LSU data.

**Table 3.**
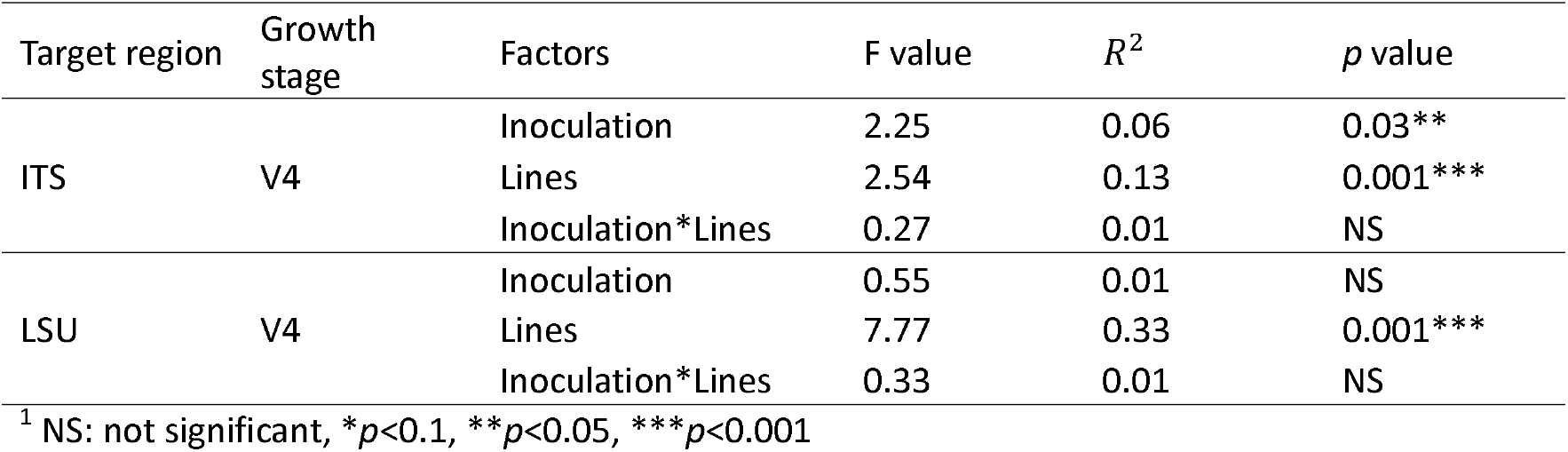
Permutation multivariate analysis of variance (PERMANOVA) to access the effect of treatments and lines at each growth stage at OTU level. The weighted Unifrac distance matrix were generated from the rarefied data set. Only growth stages with statistically significant results are listed in the table.

| Target region | Growth stage | Factors | F value | $R^2$ | $p$ value |
| --- | --- | --- | --- | --- | --- |
| ITS | V4 | Inoculation | 2.25 | 0.06 | 0.03** |
|  |  | Lines | 2.54 | 0.13 | 0.001*** |
|  |  | Inoculation*Lines | 0.27 | 0.01 | NS |
| LSU | V4 | Inoculation | 0.55 | 0.01 | NS |
|  |  | Lines | 7.77 | 0.33 | 0.001*** |
|  |  | Inoculation*Lines | 0.33 | 0.01 | NS |

<sup>1</sup> NS: not significant, \* $p < 0.1$ , \*\* $p < 0.05$ , \*\*\* $p < 0.001$

### Core microbiome

The soybean phyllosphere mycobiome was dominated by Ascomycota and accounted for 97% of classified reads. A total of 10 OTUs were identified as the core mycobiome in the soybean phyllosphere. Eight out of 10 OTUs were enriched and continuously presented across samples when using the entire data set or only the control data set (NIC/No_Fun) for analysis. OTU17 (*Septoria*, 1.6%) was identified as a core microbiome when using all samples for analysis but not in the control samples. The OTU45 (unclassified_Pleosporales, 0.14%) was identified as a core microbiome when using the LSU data set for analysis. Overall, the most abundant genus was *Gibberella* with 33.7% of classified reads, followed by *Alternaria* (24.24%), *Didymella* (11.77%), *Cladosporium* (9.02%), *Plectosphaerella* (7.67%), *Colletotrichum* (2.59%), and *Bipolaris* (2.20%) (Table 4). The core OTUs comprised 92% to 98% RA of the fungal community in each sample, and these organisms were observed across growth stages (Figure 4, Supplementary Figure 1.). The analysis of variance indicated that, at the V4 stage, the inoculation treatment had a significant effect on *Septoria* (OTU 17, *p*=0.0262), and the lines were significant for *Bipolaris* (OTU 14, *p*=0.0281). There were no significant differences detected for any core OTU at the R3 stage. At R5 stage, no significance was detected for fungicide treatments on any core OTU, but the fungicide treatment had a significant effect on *Diaporthe* (OTU11, *p*=0.0376) (Supplementary Table 8).

**Figure 4.**
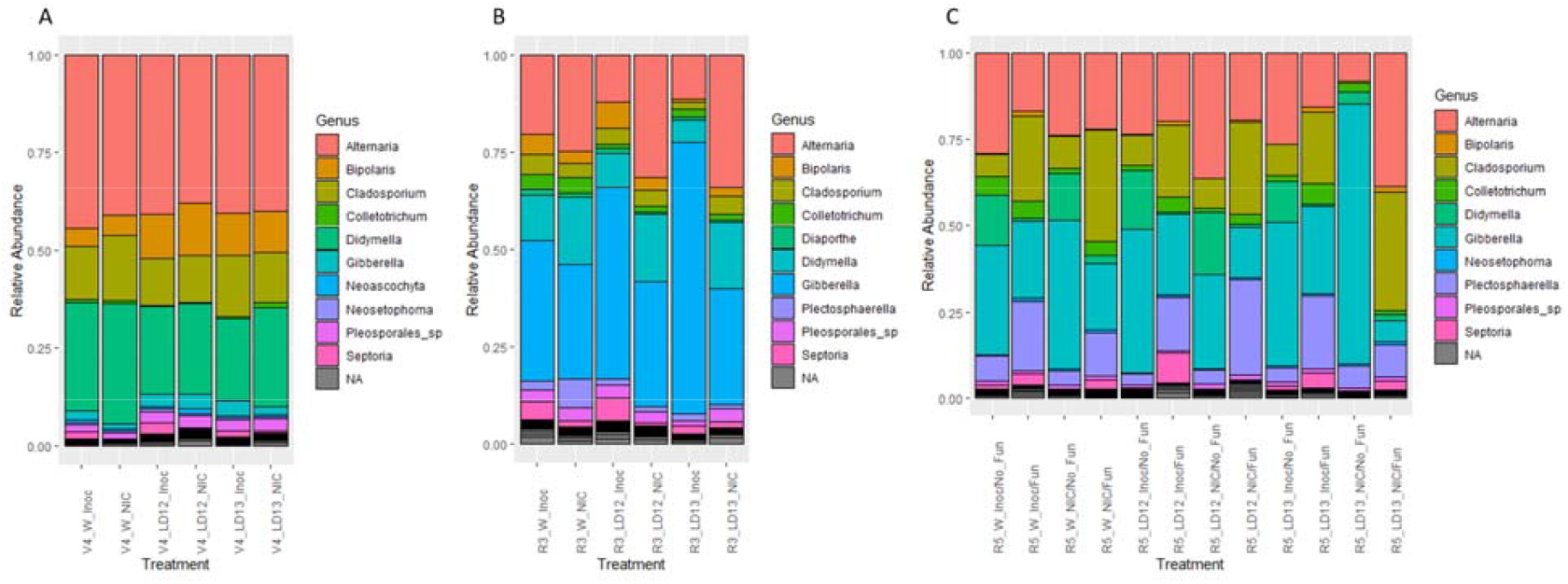
Stacked bar plots showing the relative abundance of fungal community across growth stage, lines, and treatments. (A) V4 (B) R3 and (C) R5. The top 20 most abundant taxa were listed on the legends. At V4 and R3 stage, each bar consisted of three replicates for W(illiams), LD12(−8677), and LD13(−14071R2). At R5 stage, the lines were merged for analysis since there was no significant difference between lines, and each bar consisted of 6 replicates.

**Table 4.** List of core fungal microbiome (genus level) in soybeans.

| OTU | Phylum | Class | Order | Family | Genus | Reads number | % of classified reads |
| --- | --- | --- | --- | --- | --- | --- | --- |
| OTU1 | Ascomycota | Sordariomycetes | Hypocreales | Nectriaceae | Gibberella | 899301 | 33.70 |
| OTU2 | Ascomycota | Dothideomycetes | Pleosporales | Pleosporaceae | Alternaria | 646797 | 24.24 |
| OTU7 | Ascomycota | Dothideomycetes | Pleosporales | Didymellaceae | Didymella | 314157 | 11.77 |
| OTU8 | Ascomycota | Dothideomycetes | Capnodiales | Cladosporiaceae | Cladosporium | 240713 | 9.02 |
| OTU4 | Ascomycota | Sordariomycetes | Glomerellales | Plectosphaerellaceae | Plectosphaerella | 204665 | 7.67 |
| OTU10 | Ascomycota | Sordariomycetes | Glomerellales | Glomerellaceae | Colletotrichum | 69035 | 2.59 |
| OTU14 | Ascomycota | Dothideomycetes | Pleosporales | Pleosporaceae | Bipolaris | 58795 | 2.20 |
| OTU17<sup>1</sup> | Ascomycota | Dothideomycetes | Capnodiales | Mycosphaerellaceae | Septoria | 44915 | 1.68 |
| OTU28 | Ascomycota | Dothideomycetes | Pleosporales | unidentified | Pleosporales_sp | 42599 | 1.60 |
| OTU45<sup>2</sup> | Ascomycota | Dothideomycetes | Pleosporales | unclassified_Pleosporales | unclassified_Pleosporales | 3745 | 0.14 |

<sup>1</sup> Present when using all samples as input data
<sup>2</sup> Present only in the LSU data set

**Supplementary Figure 1**. Relative abundance of core microbiomes at V4, R3, and R5 stage. At V4 and R3 stage, each box consisted of three replicates for W(illiams), LD12(−8677), and LD13(−14071R2). At R5 stage, the lines were merged for analysis since there was no significant difference between lines, and each box consisted of 6 replicates. The Wilcoxon rank-sum tests were performed for treatments (Inoc vs NIC, Fun vs No_Fun).(\**p*<0.1, \*\**p*<0.05, \*\*\**p*<0.005)

### Differential abundance of OTUs

At V4 and R3 stage, when comparing the Inoculated and non-inoculated samples, only *Septoria* (OTU17) and *Didymella* (OTU7) were significantly more abundant in the inoculated samples. Six abundance comparisons between the treatments at the R5 stage resulted in 28 and 19 significantly different OTUs using ITS (Figure 5) and LSU data sets. One of the abundance comparisons (Inoc/Fun vs Not_inoc/Fun) resulted in zero significantly different OTUs. Fifteen out of 19 OTUs with significantly different abundance on the LSU data set had consistent and similar differential patterns with the ITS data set. At the R5 stage, inoculation with *S. glycines* increased the abundance of *Septoria* (OTU17) and *Tausonia* (OTU23) when compared with non-inoculated samples (Figure 5 D). The fungicide application affected most of the core OTUs, except for *Plectosphaerella* (OTU4) and *Colletotrichum* (OTU10). Fungicide application resulted on significant decreases for OTUs such as *Epicoccum* (OTU278), *Didymella* (OTU7), and Cladosporiaceae_sp (OTU76025) (Figure 5 A and E). However, the amount of *Septoria* (OTU17), *Bipolaris* (OTU14), and *Diaporthe* (OTU11) increased on the fungicide treated samples (Figure 5 A and E, Supplementary Table 9). Differential abundance of non-targets yeasts was also identified in this analysis, for example *Hannaella* (OTU18) and *Papiliotrema* (OTU76) were significantly more abundant in the control (NIC/No_Fun) samples (Figure 5 E), and *Tausonia* (OTU 23), *Sporobolomyces* (OTU 20) and Tilletiopsis (OTU 31) significantly increased in the fungicide treated samples (Figure 5 A and B).

**Figure 5.**
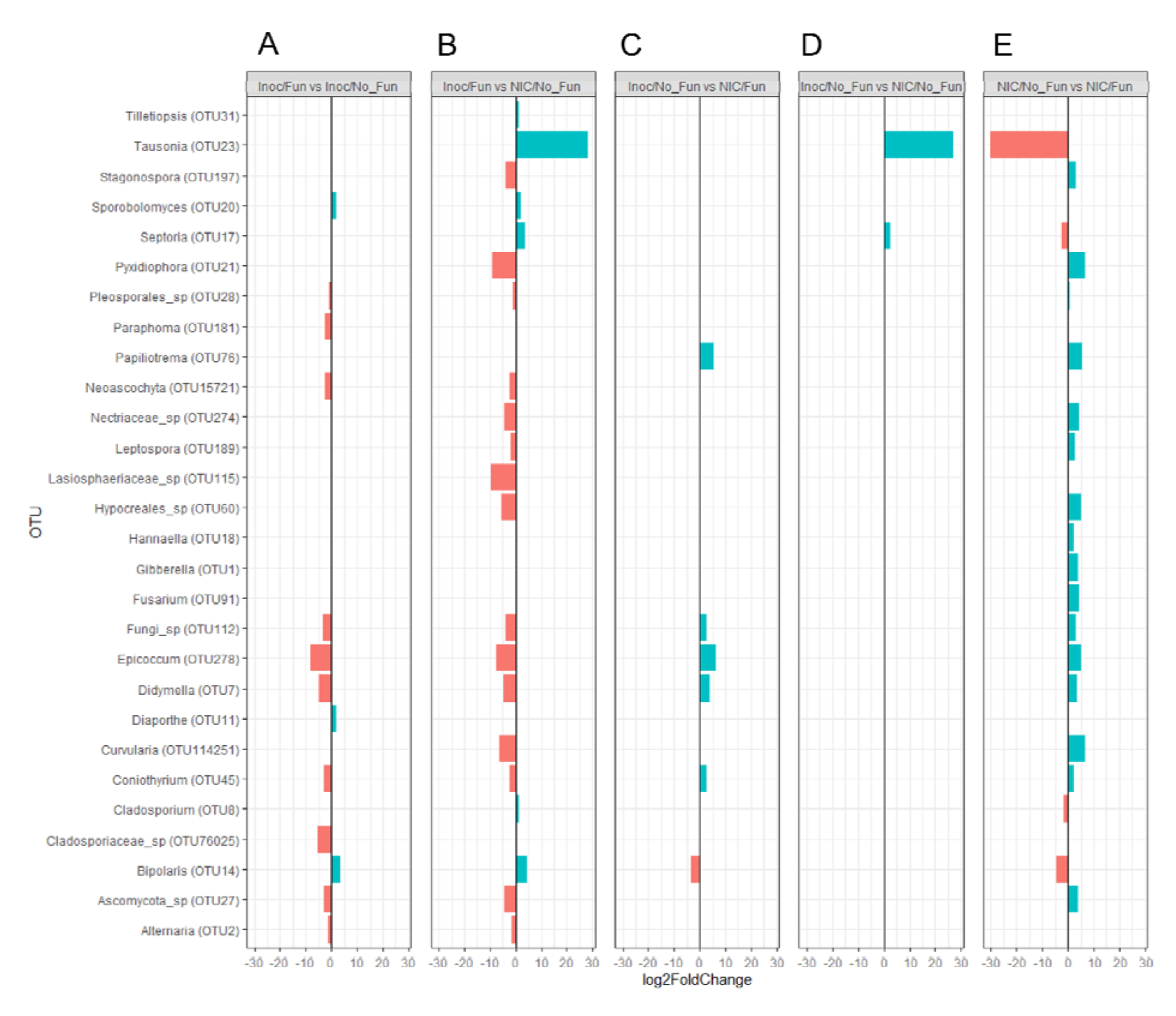
Differential abundance of OTUs between treatments at R5 stage. Shown are the significant different (FDR adjusted p-value <0.05) OTUs at the genus level with the ITS dataset. Each column (A-E) represents a comparison between treatment combinations with abbreviations: Inoc(ulated), NIC(non-inoculated), Fun(gicide), and No_Fun(gicide). A blue bar indicates a higher abundance of OTUs, and a red bar indicates a lower abundance of OTUs in the former treatment for each comparison. In (A) and (B), a blue bar indicates a higher abundance of OTUs in inoculated and fungicide treated samples; in (C) and (D), a blue bar indicates a higher abundance of OTUs in the inoculated samples; and in (E), a blue bar indicates a higher abundance of OTUs in the control samples.

### Network analysis

The data was subset by growth stages and treatments, resulting in 11 data sets for network analysis. After filtering out the low abundance OTUs, the remaining OTUs for analysis ranged between 95 to 187 (Supplementary Table 10). There were more positive edges in all networks than negative edges (Figure 6, Supplementary Figures 2-5). The networks at the later growth stage had higher numbers of nodes and edges (Supplementary Table 10, Supplementary Figure 2). The average degree and betweenness centrality increased throughout the season, indicating higher cooccurrences and bridged connections between organisms, and the complexity of the network was higher at the later soybean developmental stage (Supplementary Table 10). Microorganisms with high betweenness centrality in the network, such as *Alternaria, Bipolaris, Cladosporium, Didymella*, and *Gibberella* were also identified as the core microbiome described above (Figure 6). The consistent results between the two analysis suggests that these organisms play an important role in the community. Only a few direct associations were identified between *Septoria* and other organisms. At V4 and R3 stage, *Septoria* was predicted to have a positive association with an OTU classified as unknown. At R5 stage, after filtering by edge stability (> 0.5), *Septoria* was predicted to have positive associations with *Diaporthe, Bipolaris*, and an unidentified OTU, and negatively interacted with a OTU classified as *Hypocreales* (Figure 6). In the differential abundance analysis, *Septoria, Diaporthe*, and *Bipolaris* were shown to have similar patterns (they were all promoted after fungicide application). The results were consistent between the network analysis and differential abundance analysis.

**Figure 6.**
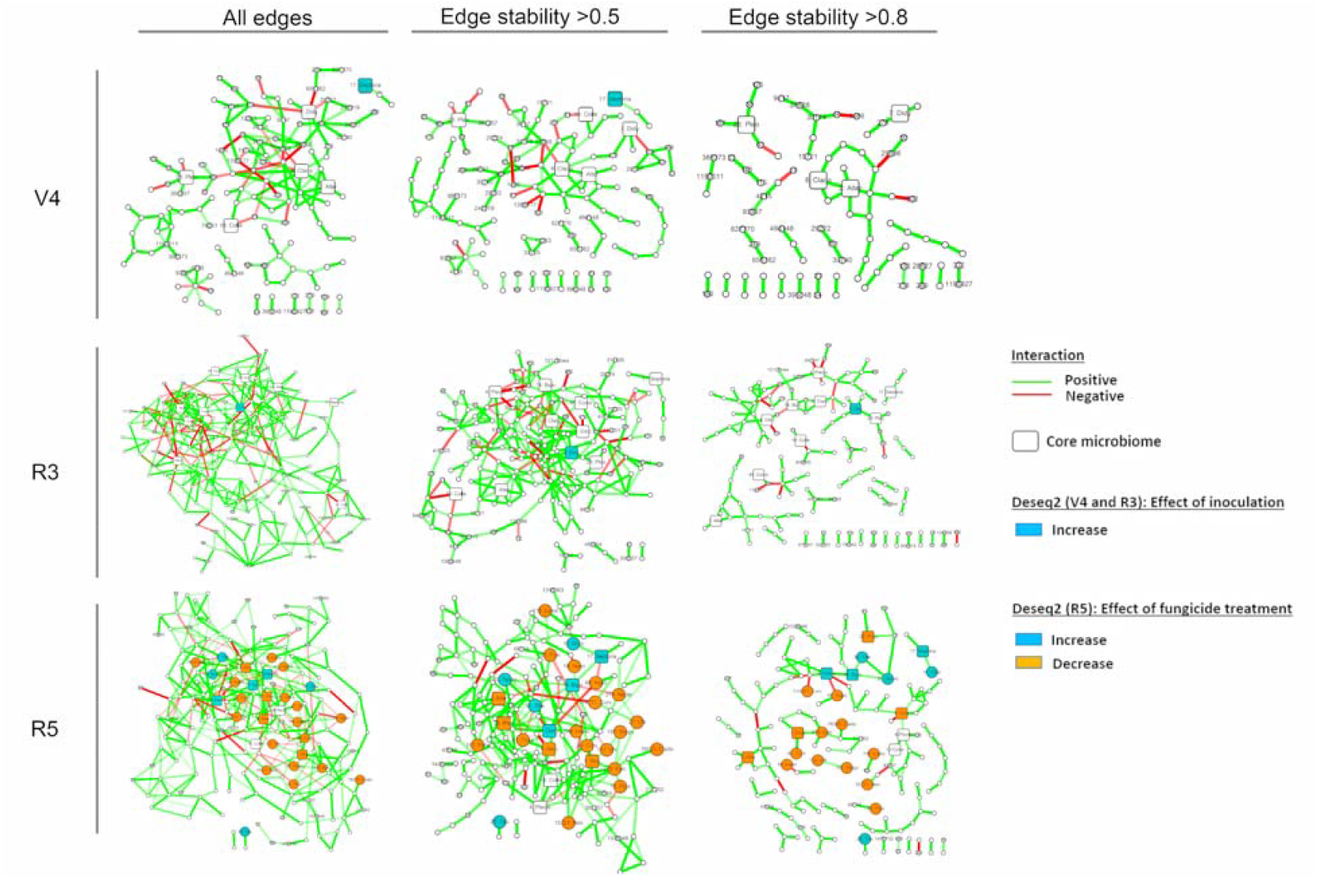
Network analysis for V4, R3, and R5 data. The networks are presented with all edges and filtered by the edge stability >0.5 and >0.8. The squared node represents the OTUs that were identified as core microbiome from the core microbiome analysis. The node color corresponds to the results of OTU differential analysis using Deseq2. Edges color corresponds to potential positive (green) and negative (red) associations between OTUs. The nodes plotted with OTU numbers were classified to a known organism at the Genus level. The nodes without labels were unclassified OTUs. The OTUs that were differentially abundant from the Deseq2 results were labeled with OTU numbers followed by the abbreviations of the genus name. Alter(naria), Bipo(laris), Clado(sporiaceae), Clado(sporium), Colle(totrichum), Coni(othyrium), Curv(ularia), Diap(orthe), Didy(mella), Epic(occum), Fusa(rium), Gibb(erella), Hann(aella), Hypo(creales), Lasio(sphaeriaceae), Lepto(spora), Nectri(aceae), Neo(ascochyta), Papi(liotrema), Para(phoma), Pleo(sporales), Pyxi(diophora), Septoria, Sporo(bolomyces), Stago(nospora), Tau(sonia), and Tille(tiopsis).

**Supplementary Figure 2 Networks for (A) V4, (B) R3 and (C) R5 data**. The squared node represents that the OTUs were identified as core microbiome from the core microbiome analysis. The node color corresponds to the OTUs at the Class level. Edges color corresponds to potential positive (green) and negative (red) interactions between OTUs.

**Supplementary Figure 3 Networks for V4 stage data (A) Inoculated and (B) non-inoculated**.

The squared node represents that the OTUs were identified as core microbiome from the core microbiome analysis. The node color corresponds to the OTUs at the Class level. Edges color corresponds to potential positive (green) and negative (red) interactions between OTUs.

**Supplementary Figure 4 Networks for R3 stage data (A) Inoculated and (B) non-inoculated**.

The squared node represents that the OTUs were identified as core microbiome from the core microbiome analysis. The node color corresponds to the OTUs at the Class level. Edges color corresponds to potential positive (green) and negative (red) interactions between OTUs.

**Supplementary Figure 5 Networks for R5 stage data (A) R5_Fun (B) R5_noFun (C) R5_Inoc, and (D) R5_NIC**. The squared node represents that the OTUs were identified as core microbiome from the core microbiome analysis. The node color corresponds to the OTUs at the Class level. Edges color corresponds to potential positive (green) and negative (red) interactions between OTUs.

## Discussion

In this study, we assessed the effect of Septoria brown spot and fungicide application on the dynamics of phyllosphere fungal communities at three soybean developmental stages: V4 (after inoculation), R3 (before fungicide application), and R5 (after fungicide application). We sequenced 96 samples using oxford nanopore technologies, and after sequence analysis obtained 3,342 OTUs or 649 genera. We identified the OTUs and genera that significantly increased or decreased after the inoculation or fungicide application treatments. The potential interactions between those organisms were estimated using network analysis. We also characterized the core mycobiome associated with soybean phyllosphere.

The phyllosphere mycobiome were significantly associated with the soybean lines only at the early growth stage. The difference between lines were not detected from the R3 and R5 samples. Sapkota et al. (2015) also observed the fungal community clustered by the variety in the non-fungicide treated samples in cereals. Also, an association between plant genotype and phyllosphere microbiome has been reported in grape species (Singh et al. 2019). However, they only collected the samples one time each year. It is unclear if this association persists throughout growth stages. More evidence suggested that the microbiome communities in the rhizosphere were associated with plant genotypes in many crops, such as maize, wheat, pea, and oat (Turner et al. 2013). In soybeans, the genotypes had a significant but mild effect on the rhizosphere microbiome when comparing five soybean genotypes, including cv. Williams, a non-nodulating mutant of Williams, cv. Williams 82, a drought-resistant cultivar, and a cyst nematode-resistance line (Liu et al. 2019). In these soybean lines, cv. Williams, cv. Williams 82, and the mutant of Williams shared similar genetic backgrounds, and they clustered together on the beta diversity analysis (Liu et al. 2019). The soybean lines we used in our study have different genetic backgrounds. Our results show that the plant genotypes influenced the microbial community in the phyllosphere at the early growth stage.

As expected, inoculation with *S. glycines* resulted on significantly higher levels of Septoria DNA amplicons. An assumed higher *S. glycines* biomass did not significantly alter the species richness and evenness (alpha diversity) of the mycobiome but some significant differences were observed for the composition of the species (beta diversity) between inoculated and not inoculated samples. The network analysis identified significant positive associations between *Septoria* reads with *Diaporthe, Bipolaris* and an unidentified OTU, while there was a negative interaction with an OTU classified as Hypocreales. Carmona et al. (2011) suggested an association between *S. glycines, Cercospora spp*. and *Colletotrichum spp*. While we did not find this same associations, the association with *Diaporthe* shows the co-occurrence of pathogens that generally cause disease at the end of the season. Differences on what pathogens affect soybean at the end of the season might be due to host genotype, environment, and geography. Our results give some support to the existence of a late season disease complex of soybean of which *S. glycines* is an important component.

Priaxor (14.3% fluxapyroxad and 28.6% pyraclostrobin) is a broad-spectrum foliar fungicide recommended to applied between R2 to R4 developmental stages targeting multiple foliar diseases, including Alternaria leaf spot, Anthracnose, Septoria brown spot, Cercospora leaf blight, Frogeye leaf spot, Pod and stem blight, and Rhizoctonia. It is reasonable to expect that fungicide application will reduce the richness and other alpha diversity indices, but we did not see these results in the R5 stage samples. However, a trend of reduced variation within fungicide treated samples than the untreated samples were observed in other studies. Both, Knorr et al. (2019) and Karlsson et al. (2014) reported higher variation within untreated samples than fungicide treated samples in fungal phyllosphere in wheat. According to our PERMANOVA analysis and PCoA plots, the fungicide treatment influenced the fungal communities on soybean leaves. The relative abundance analysis suggested that the fungicide application significantly reduced the amount of *Alternaria, Epicoccum, Didymella*, and others. However, it also significantly increased the amount of *Bipolaris* and *Diaporthe*. Some of the patterns we observed were consistent with the results in Batzer and Mueller (2020). Both results highlight the possibility of increasing the biomass of some pathogenic organisms with the application of fungicide.

Surprisingly, the fungicide application did not influence the relative abundance of the *Septoria* OTU at the R5 stage samples. However, this non-significant fungicide effect on target pathogens was also observed in other phyllosphere mycobiome studies in small grains (Karlsson et al. 2014; Sapkota et al. 2015). Karlsson et al 2014, reported that the fungicide did not significantly affect common wheat pathogens identified from their study, such as *Mycosphaerella graminicola, Blumeria graminis, Puccinia striiformis, Phaeosphaeria nodorum* (*Parastagonospora nodorum*), *Monographella spp*, and *Pyrenophora tritici-repentis*. Sapkota et al. 2015 also reported that the fungicide did not have a significant effect on the fungicide target organisms, *Zymoseptoria tritici* and *Blumeria graminis*. One of the possible reasons mentioned in Sapkota et al (2015) was the fast resilience of the fungal phyllosphere communities after fungicide application. The time from fungicide application to sample collection might enable the pathogens to recover. In our case, when considering the characteristics of the *S. glycines*, the pathogen first colonizes the lower canopy and gradually develops to the upper canopy. While we apply fungicide from the top canopy, it is possible that the pathogen escaped from the application, and because the fungicide reduced the organisms that occupied the space in the upper canopy it allowed Septoria to easily colonize the now available tissue.

The core microbiome has been defined as a set of stable and persistent microorganisms associated with specific status or environmental conditions (Berg et al. 2020). In this study, a total of 8 OTUs were identified as core microbiome. Many of those organisms (*Alternaria, Cladosporium, Colletotrichum, Fusarium*, and *Didymella*) have been identified as endophytes in soybeans using cultural-dependent methods (Impullitti and Malvick 2013; Batzer and Mueller 2020). In our study, although the sequences were consistently identified and enriched across all samples, a dynamic composition of the core microbes between growth stage and treatments was observed. Lemanceau et al (2017) have reviewed examples of functional core microbiota (mostly bacteria) that can enhance host fitness regardless of plant nutrient intake, drought tolerance, and defense response. The function of the core microbes on soybeans is still unclear and need further characterization. Our study showed that the soybean fungal phyllosphere was dominated by Ascomycota with 97% of classified reads. However, Noel et al (2021) reported a high number of Basidiomycyete yeasts in the soybean phyllosphere, which was not identified as core microbiome in our study. This is possibly due to the choice of primers for sequencing and databases (eukaryote database or fungal database) for taxonomy assigiment.

This study used oxford nanopore sequencing technology to generate long amplicons covering the full-length ITS and partial LSU regions to study the phyllosphere microbiome in soybeans. We highlight the effect of lines and fungicide treatment on the fungal communities on soybean leaves. We also investigated the potential association between *Septoria* and other fungal organisms. Although we are still unable to identify OTUs to the species level due to limitations of the database, this may be resolved in the near future when third-generation sequencing technologies applied to more studies, and more full-length ITS and LSU sequences become available in the database. We also believed that understanding the dynamic of the microbiome in the phyllosphere can facilitate the strategies for disease management. Our results reveal the soybean phyllosphere mycobiome and open the possibilities of targeted manipulation of fungal infection to improve foliar disease management.

## Acknowledgements

This work was supported by USDA National Institute of Food and Agriculture, Hatch project ILLU-802-922 / Accession No. 1008676. We thank Dr. Anthony Yannarell and Dr. Maria B. Villamil for the guidance, Juan P. Granda for technical assistance, and Jessica Holmes for training on data analysis.

