## Supplementary Figures for "The effect of *Septoria glycines* and fungicide application on the soybean phyllosphere mycobiome"

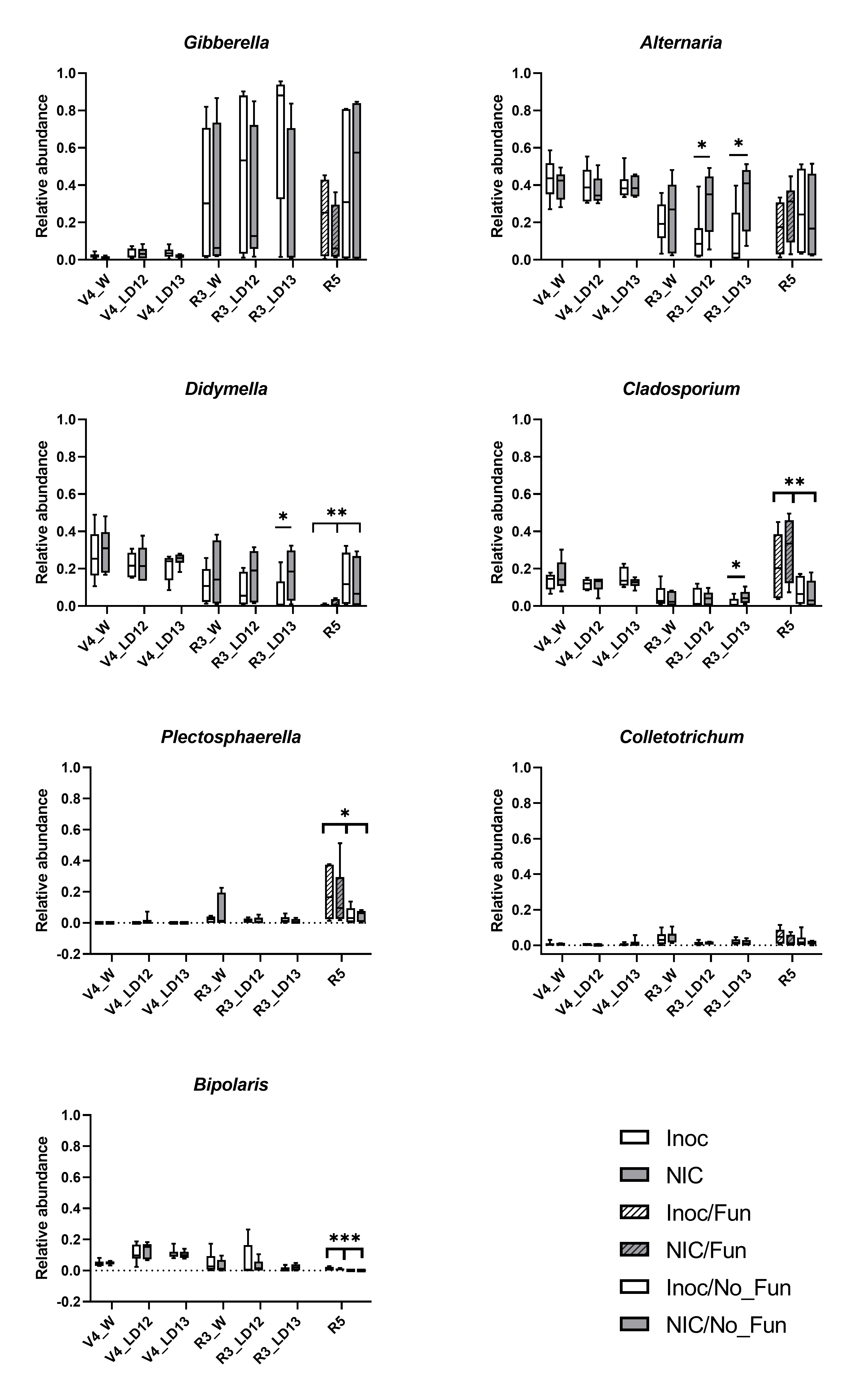

**Supplementary Figure 1.** Relative abundance of core microbiomes at V4, R3, and R5 stage. At V4 and R3 stage, each box consisted of three replicates for W(illiams), LD12(-8677), and LD13(-14071R2). At R5 stage, the lines were merged for analysis since there was no significant difference between lines, and each box consisted of 6 replicates. The Wilcoxon rank-sum tests were performed for treatments (Inoc vs NIC, Fun vs No_Fun).( *p<0.1, **p<0.05, ***p<0.005)

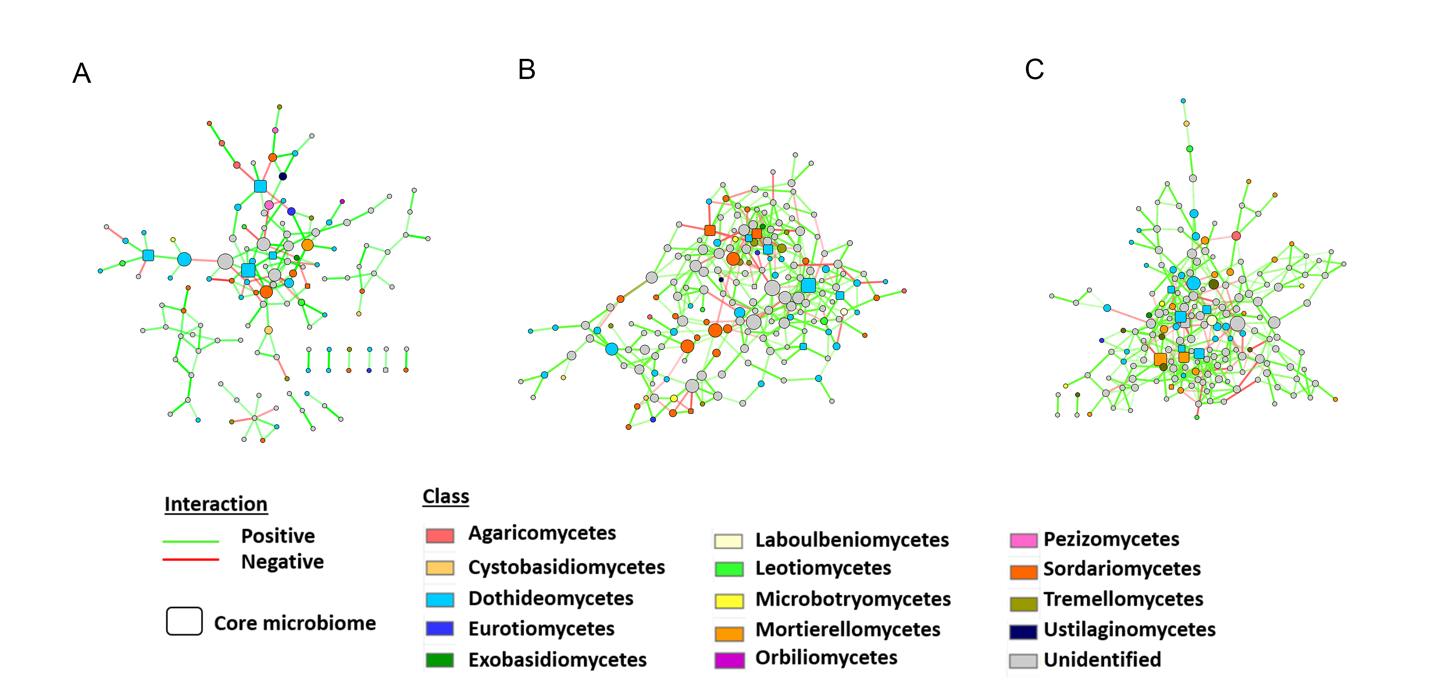

**Supplementary Figure 2** Networks for (A) V4, (B) R3 and (C) R5 data. The squared node represents that the OTUs were identified as core microbiome from the core microbiome analysis. The node color corresponds to the OTUs at the Class level. Edges color corresponds to potential positive (green) and negative (red) interactions between OTUs.

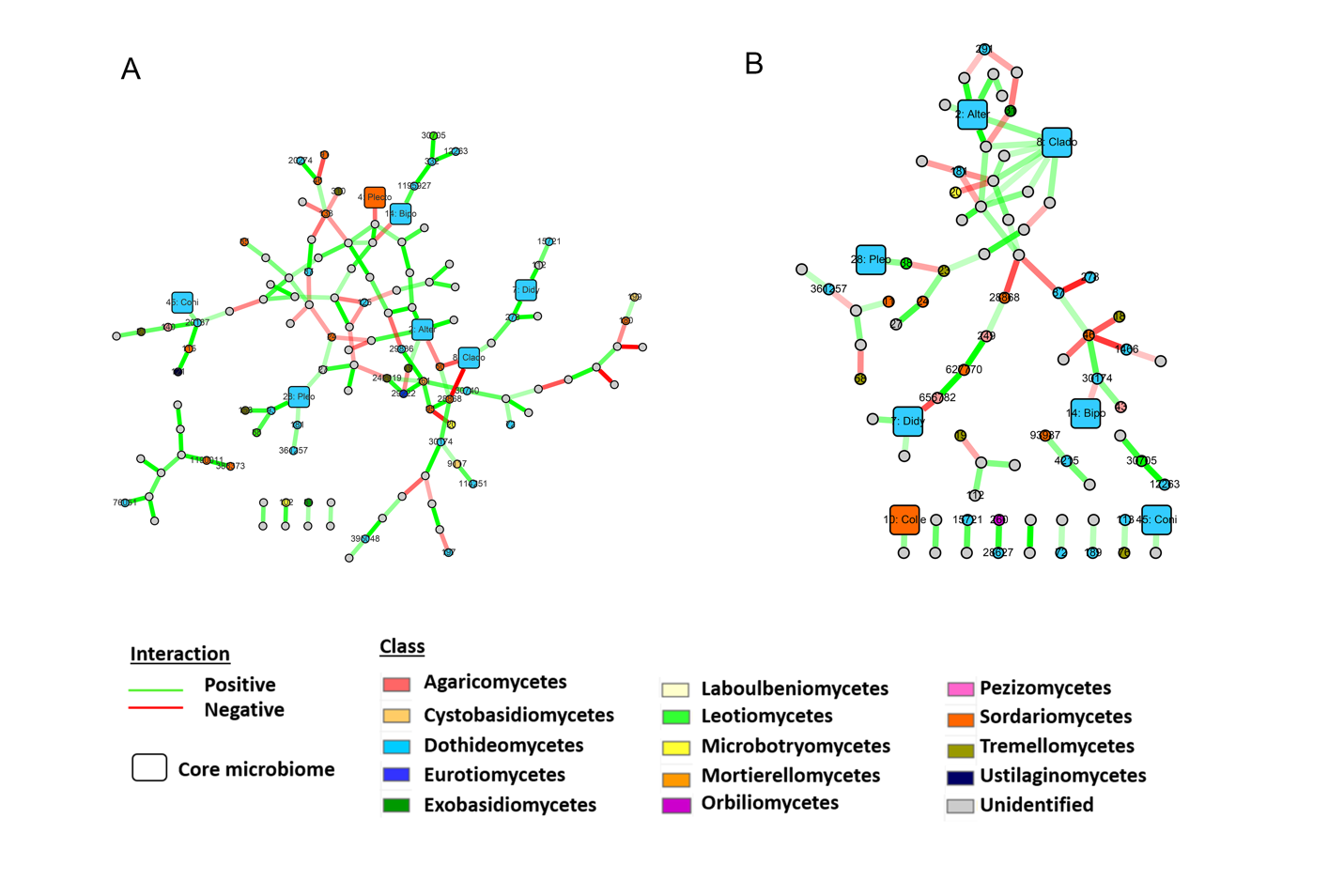

**Supplementary Figure 3 Networks for V4 stage data (A) Inoculated and (B) non-inoculated.** The squared node represents that the OTUs were identified as core microbiome from the core microbiome analysis. The node color corresponds to the OTUs at the Class level. Edges color corresponds to potential positive (green) and negative (red) interactions between OTUs.

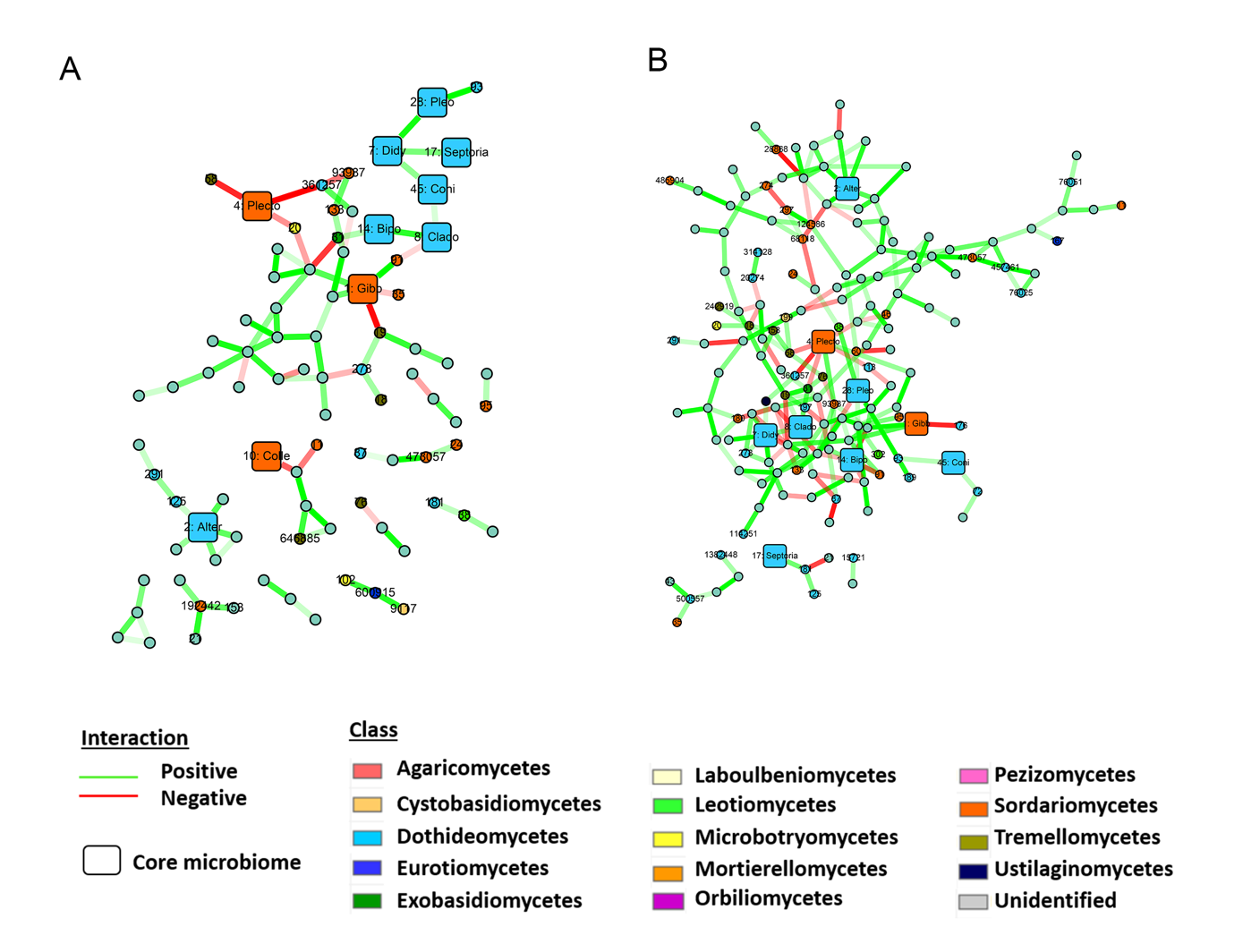

**Supplementary Figure 4** Networks for R3 stage data (A) Inoculated and (B) non-inoculated. The squared node represents that the OTUs were identified as core microbiome from the core microbiome analysis. The node color corresponds to the OTUs at the Class level. Edges color corresponds to potential positive (green) and negative (red) interactions between OTUs.

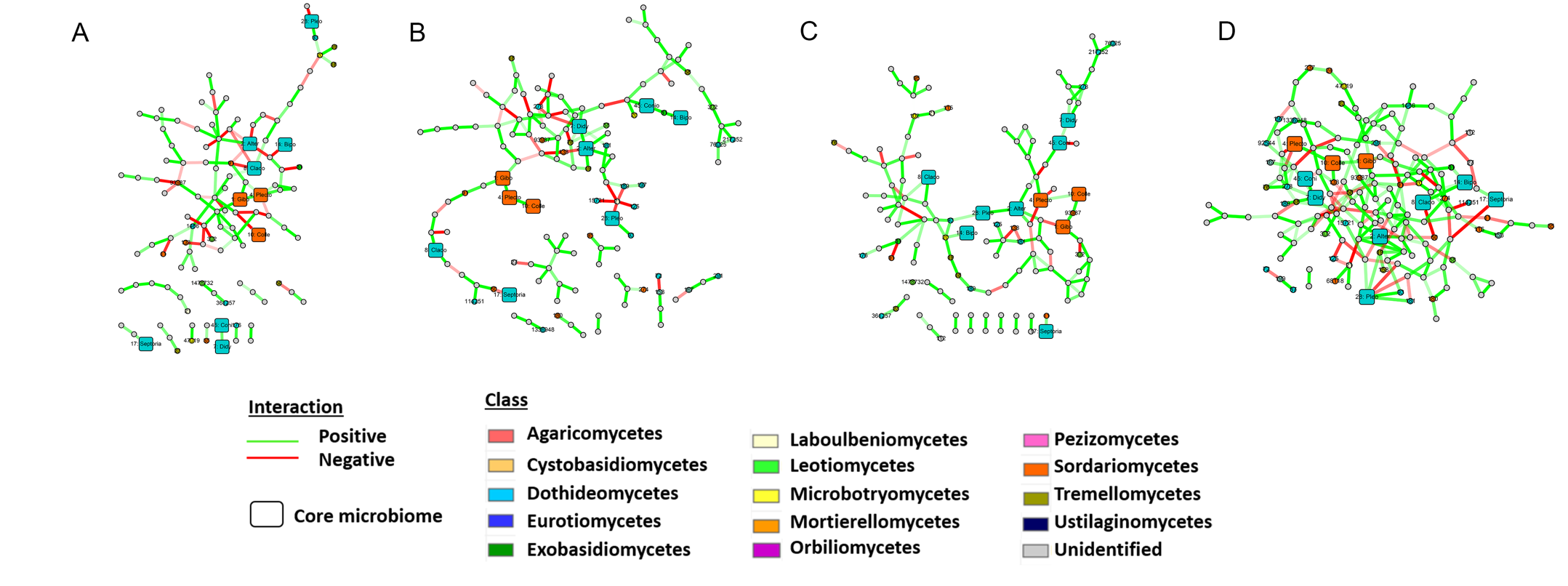

**Supplementary Figure 5** Networks for R5 stage data (A) R5_Fun (B) R5_noFun (C) R5_Inoc, and (D) R5_NIC . The squared node represents that the OTUs were identified as core microbiome from the core microbiome analysis. The node color corresponds to the OTUs at the Class level. Edges color corresponds to potential positive (green) and negative (red) interactions between OTUs.
